# High Activity of Selective Essential Oils against Stationary Phase *Borrelia burgdorferi*

**DOI:** 10.1101/130898

**Authors:** Jie Feng, Shuo Zhang, Wanliang Shi, Nevena Zubcevik, Judith Miklossy, Ying Zhang

## Abstract

Although the majority of patients with Lyme disease can be cured with the standard 2-4 week antibiotic treatment, about 10-20% of patients continue to suffer from post-treatment Lyme disease syndrome (PTLDS). While the cause for this is debated, one possibility is due to persisters not killed by the current Lyme antibiotics. It has been reported that essential oils have antimicrobial activities and some have been used by patients with persisting Lyme disease symptoms. However, the activity of essential oils against the causative agent *Borrelia burgdorferi (B. burgdorferi)* has not been carefully studied. Here, we evaluated the activity of 34 essential oils against *B. burgdorferi* stationary phase culture as a model for persisters. We found that many essential oils had varying degrees of activity against *B. burgdorferi*, with top 5 essential oils (oregano, cinnamon bark, clove bud, citronella, and wintergreen) at a low concentration of 0.25% showing more activity than the persister drug daptomycin. Interestingly, some highly active essential oils were found to have excellent anti-biofilm ability as shown by their ability to dissolve the aggregated biofilm-like structures. The top 3 hits, oregano, cinnamon bark and clove bud, completely eradicated all viable cells without regrowth in subculture. Carvacrol was found to be the most active ingredient of oregano oil showing excellent activity against *B. burgdorferi* stationary phase cells, while p-cymene and α-terpinene had no apparent activity. Future studies are needed to characterize and optimize the active essential oils in drug combinations in vitro and in vivo for improved treatment of persistent Lyme disease.

**IMPORTANCE:** There is a huge need for effective treatment of patients with Lyme disease who suffer from PTLDS. Recent in vitro and in vivo studies suggest that *B. burgdorferi* develops persisters that are not killed by the current Lyme antibiotics as a possible contributor to this condition. Although essential oils are used by patients with Lyme disease with variable improvement in symptoms, their anti-borrelia activity has not been carefully studied. Here we found that not all essential oils have adequate anti-borrelia activity and identified some highly potent essential oils (oregano, cinnamon bark, clove bud) that have even higher anti-persister and anti-biofilm activity than the persister drug daptomycin. Carvacrol was found to be the most active ingredient of oregano oil and have the potential to serve as a promising oral persister drug. Our findings may have implications for developing improved treatment of persisting Lyme disease.

## INTRODUCTION

Lyme disease, which is caused by *Borrelia burgdorferi (B. burgdorferi)* sensu lato complex species, is the most common vector-borne disease in the United States with an estimated 300,000 cases a year (1). The infection is transmitted to humans by tick vectors that feed upon rodents, reptiles, birds, and deer, etc. (2). In the early stage of Lyme disease, patients often have localized erythema migrans rash that expands as the bacteria disseminate from the cutaneous infection site via blood stream to other parts of the body. Late stage Lyme disease is a multi-system disorder which can cause arthritis and neurologic manifestations (1). While the majority of Lyme disease patients can be cured if treated early with the standard 2-4 week doxycycline, amoxicillin, or cefuroxime therapy (3), at least 10-20% of patients with Lyme disease have lingering symptoms such as fatigue, muscular and joint pain, and neurologic impairment even 6 months after the antibiotic treatment - a set of symptoms called Post-Treatment Lyme Disease Syndrome (PTLDS) (4). While the cause of PTLDS is unknown, several possibilities may be involved, including autoimmune response (5), immune response to continued presence of antigenic debris (6), tissue damage as a result of Borrelia infection and inflammation, co-infections (7), as well as persistent infection due to *B. burgdorferi* persisters that are not killed by the current antibiotics used to treat Lyme disease (8–10). Various studies have found evidence of *B. burgdorferi* persistence in dogs (11), mice (8, 9), monkeys (10), as well as humans (12) after antibiotic treatment, however, viable organisms are very difficult to be cultured from the host after antibiotic treatment.

In log phase cultures (3-5 day old), *B. burgdorferi* is primarily in motile spirochetal form which is highly susceptible to current Lyme antibiotics doxycycline and amoxicillin, however, in stationary phase cultures (7-15 day old), increased numbers of atypical variant forms such as round bodies and aggregated biofilm-like microcolonies develop (13, 14). These atypical forms have increased tolerance to doxycycline and amoxicillin when compared to the growing spirochetal forms (13–16). In addition, that the active hits from the round body persister screens (17) overlap with those from the screens on stationary phase cells (13) indicates the stationary phase culture contains overlapping persister population and can be used as a relevant persister model for drug screens to identify agents with anti-persister activity. Using these models,we identified a range of drugs such as daptomycin, clofazimine, anthracycline antibiotics, and sulfa drugs with high activity against stationary phase cells enriched in persisters through screens of FDA-approved drug library and NCI compound libraries (13, 18).

Essential oils are concentrated volatile liquid that are extracted from plants. It has been reported in the literature that essential oils have antimicrobial activities (19) and anecdotal reports from the internet suggest some essential oils may improve symptoms for patients with persistent Lyme disease symptoms. However, the activity of essential oils against the causative agent *B. burgdorferi* has not been properly studied. Here, we evaluated a panel of essential oils for activities against *B. burgdorferi* stationary phase cells, and found that not all essential oils used by patients with Lyme disease have the same activity against *B. burgdorferi*, with oregano, cinnamon bark, and clove bud having among the highest anti-persister activity in vitro.

## RESULTS

### Evaluation of essential oils for activity against stationary phase *B. burgdorferi*

We evaluated a panel of 34 essential oils at four different concentrations (1%, 0.5%, 0.25% and 0.125%) for activity against a 7-day old *B. burgdorferi* stationary phase culture in the 96-well plates with control drugs for 7 days. Consistent with our previous studies (13, 20), daptomycin control was shown to have high activity against the *B. burgdorferi* stationary phase culture, with a dose-dependent increase in killing activity resulting in a near total clearance of *B. burgdorferi* cells at the 40 μM concentration (Figure 1). Five essential oils (bandit, oregano, clove bud, geranium bourbon and cinnamon bark) at 1% concentration showed more activity against the stationary phase *B. burgdorferi* culture than 40 μM daptomycin with the plate reader SYBR green I/PI assay (Table 1). We found some essential oils have autofluorescence which severely interfered with the SYBR Green I/PI plate reader assay, but we were able to identify and resolve this issue present in some samples by fluorescence microscopy. As we previously described (21), we directly calculated the green (live) cell ratio of microscope images using Image Pro-Plus software, which could eliminate the background autofluorescence. Using SYBR Green I/PI assay and fluorescence microscopy, we additionally found 18 essential oils that showed more or similar activity against the stationary phase *B. burgdorferi* at 1% concentration compared to the 40 μM daptomycin, which could eradicate all live cells as shown by red (dead) aggregated cells (Table 1; Figure 1A). At 0.5% concentration, 7 essential oils (oregano, cinnamon bark, clove bud, citronella, wintergreen, geranium bourbon, and patchouli dark) were found to have higher or similar activity against the stationary phase *B. burgdorferi* than 40 μM daptomycin by fluorescence microscope counting after SYBR Green I/PI assay (Table 1; Figure 1B). However, bandit thieves oil, while having good activity at 1%, had significantly less activity at 0.5% and lower concentrations (Table 1). Among the effective hits, 5 essential oils (oregano, cinnamon bark, clove bud, citronella, and wintergreen) still showed better activity than 40 μM daptomycin at 0.25% concentration (Table 1; Figure 1C). Eventually, oregano, cinnamon bark, and clove bud were identified as the most active essential oils because of their remarkable activity even at the lowest concentration of 0.125%, which showed similar or better activity than 40 μM daptomycin (Table 1; Figure 1D).

**FIG 1.**
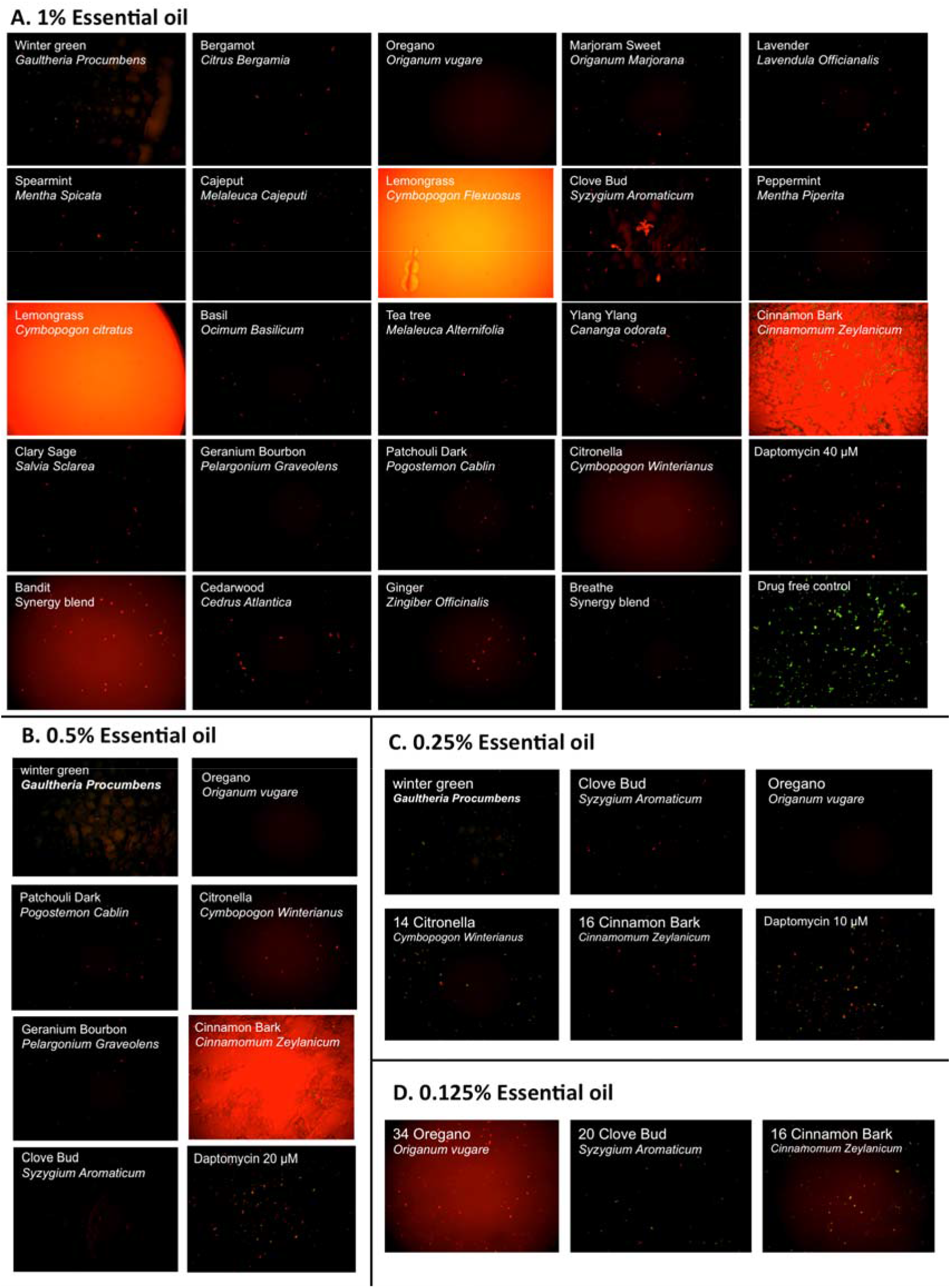
Effect of essential oils on the viability of stationary phase *B. burgdorferi*. A 7- day old *B. burgdorferi* stationary phase culture was treated with essential oils at different concentrations (v/v), 1% (A), 0.5% (B), 0.25% (C), and 0.125% (D) for 7 days followed by staining with SYBR Green I/PI viability assay and fluorescence microscopy.

**Table 1.**
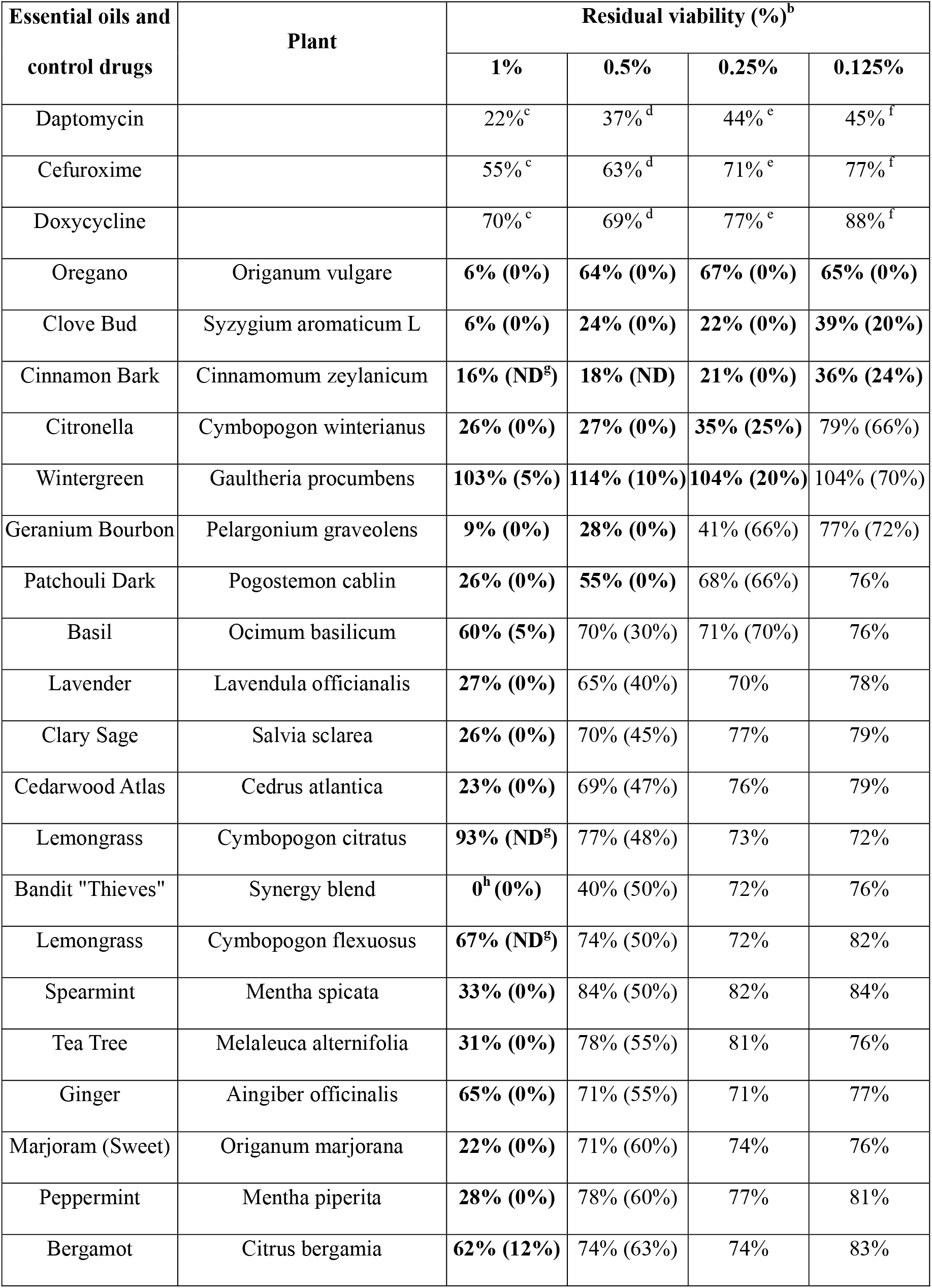
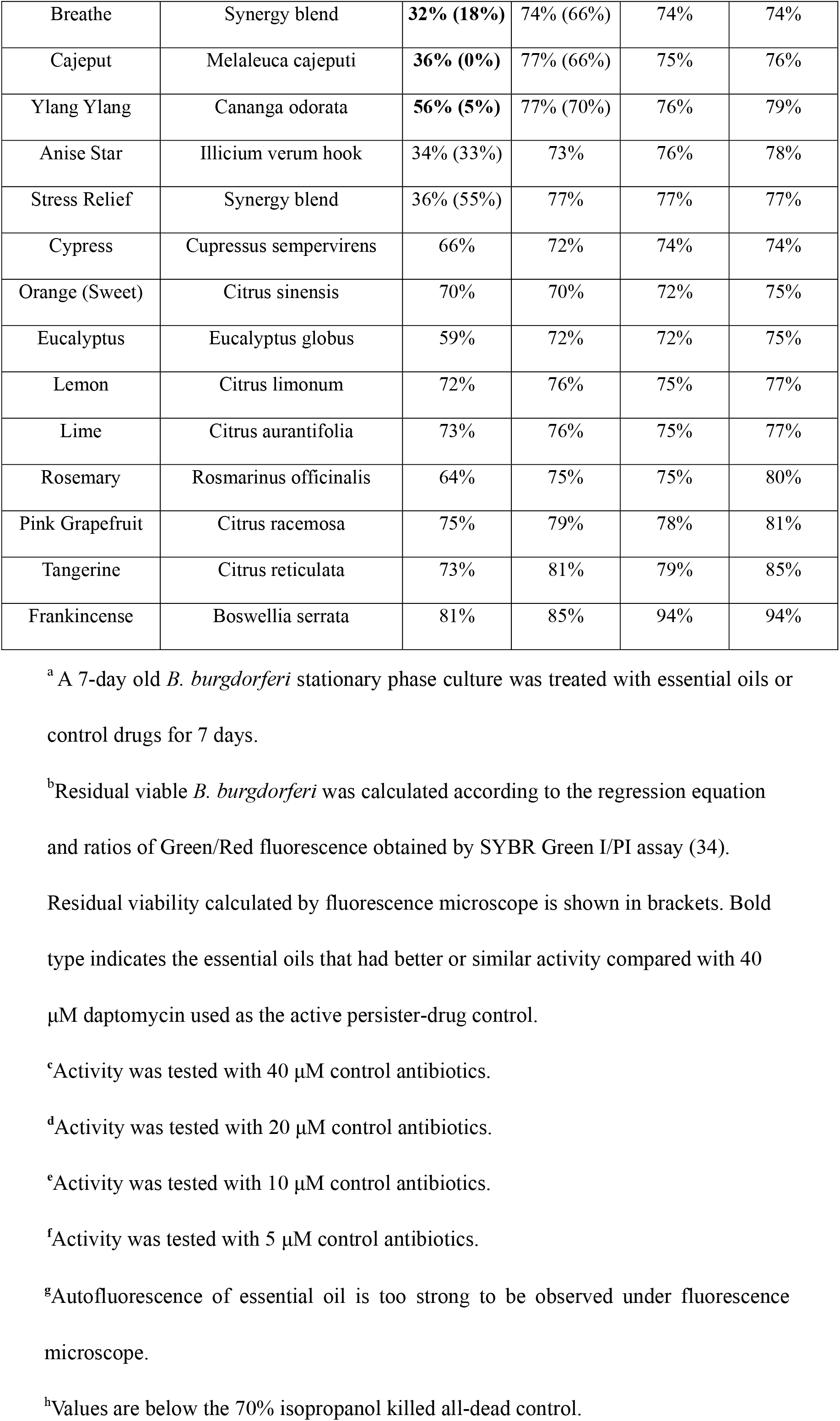
Effect of essential oils on a 7-day old stationary phase *B. burgdorferi*^a^.

To further compare the activity of these active essential oils and find whether they could eradicate stationary phase *B. burgdorferi* at lower concentrations, we evaluated 6 essential oils (oregano, cinnamon bark, clove bud, citronella, geranium bourbon, and wintergreen) at even lower concentrations at 0.1% and 0.05%. We noticed that oregano could not wipe out stationary phase *B. burgdorferi* at 0.05% concentration as shown by some residual green aggregated cells (Table 2, Figure 2), despite oregano showed strong activity sterilizing all the stationary phase *B. burgdorferi* cells at above 0.1% concentration (Tables 1 and 2).

**FIG 2.**
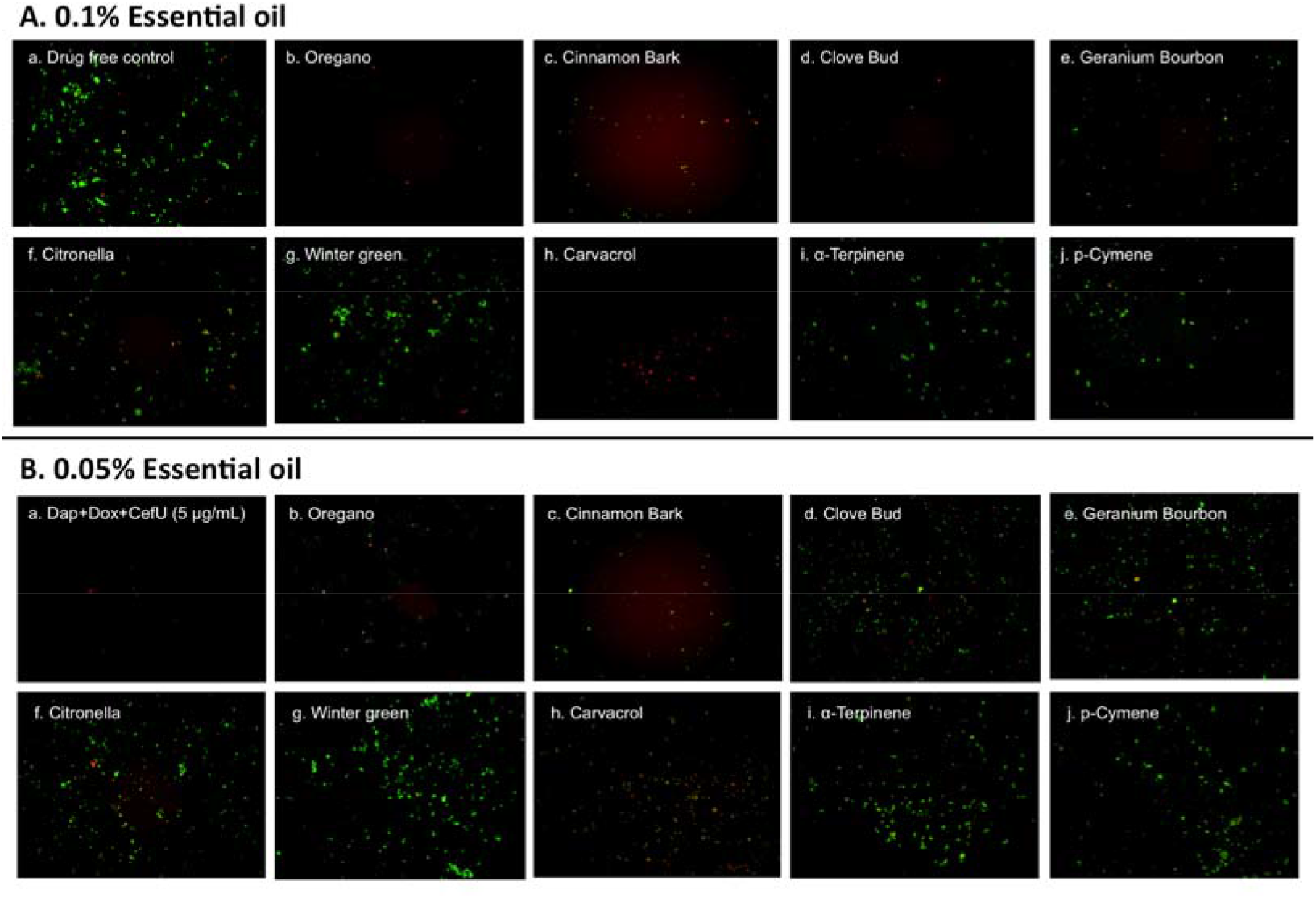
Effect of active essential oils or their ingredients on stationary phase *B. burgdorferi*. A *B. burgdorferi* stationary phase culture (7-day old) was treated with 0.1% (A) or 0.05% (B) essential oils (labeled on the image) or the ingredients (carvacrol, α–terpinene or p–cymene) of oregano for 7 days followed by staining with SYBR Green I/PI viability assay and fluorescence microscopy.

**Table 2.** Comparison of essential oil activity against stationary phase *B. burgdorferi* with 0.1% and 0.05% (v/v) treatment and subculture^a^.

|  | 0.1% Essential oil |  | 0.05% Essential oil |  |
| --- | --- | --- | --- | --- |
|  | Treatment<sup>b</sup> | Subculture<sup>c</sup> | Treatment | Subculture<sup>c</sup> |
| Drug free control | 95% | + | 95% | + |
| Daptomycin+Doxycycline+Cefuroxime<sup>d</sup> | 18%<sup>d</sup> | -<sup>d</sup> | N/A | N/A |
| Oregano | 60% (8%) | - | 68% (56%) | - |
| Cinnamon Bark | 62% (55%) | - | 66% (66%) | - |
| Clove Bud | 57% (33%) | - | 68% (77%) | + |
| Citronella | 78% (70%) | + | 77% (82%) | + |
| Geranium Bourbon | 74% (70%) | + | 85% (80%) | + |
| Wintergreen | 90% (77%) | + | 94% (85%) | + |
| Carvacrol | 55% (2%) | - | 60% (55%) | - |
| p-cymene | 66% (72%) | + | 73% (83%) | + |
| α-terpinene | 70% (77%) | + | 77% (85%) | + |

<sup>a</sup> A 7-day old stationary phase *B. burgdorferi* was treated with 0.05% or 0.1 % essential oils or their ingredients for 7 days when the viability of the residual organisms was assessed by subculture.
<sup>b</sup>Residual viable percentage of *B. burgdorferi* was calculated according to the regression equation and ratio of Green/Red fluorescence obtained by SYBR Green I/PI assay as described (13). Direct microscopy counting was employed to rectify the results of the SYBR Green I/PI assay. Residual viability calculated by fluorescence microscopy is shown in brackets. Viabilities are the average of three replicates.
<sup>c</sup>“+” indicates growth in subculture; “-” indicates no growth in subculture.
<sup>d</sup>Activity was tested with 5 µg/mL antibiotic combination.

### Carvacrol as a highly potent active ingredient of oregano oil against stationary phase

*B. burgdorferi*. To identify active ingredients of the oregano essential oil, we tested three major constituents (22), carvacrol, p-cymene and α-terpinene on the stationary phase *B. burgdorferi*. Interestingly, carvacrol showed similar high activity against *B. burgdorferi* as oregano essential oil either at 0.1% (6.5 μM) or 0.05% (3.2 μM) concentration (Table 2 and Figure 2h). Meanwhile we also found carvacrol was very active against replicating *B. burgdorferi*, as shown with a very low MIC of 0.16-0.31 μg/mL. By contrast, p-cymene and α-terpinene did not have activity against the stationary phase *B. burgdorferi* (Table 2 and Figure 2i and j). Thus, carvacrol could be one of the most active ingredients in oregano oil that kill stationary phase *B. burgdorferi*.

### Subculture studies to evaluate the activity of essential oils against stationary phase *B. burgdorferi*

To confirm the activity of the essential oils in killing stationary phase *B. burgdorferi*, we performed subculture studies in BSK-H medium as described previously (14). To validate the activity of these essential oils, samples of essential oil treated cultures were subjected to subculture after removal of the drugs by washing followed by incubation in fresh BSK medium for 21 days. According to the essential oil drug exposure experiments (Table 2), we used subculture to further confirm whether the top 6 active essential oils (oregano, cinnamon bark and clove bud, citronella, geranium bourbon, and wintergreen) could eradicate the stationary phase *B. burgdorferi* cells at 0.1% or 0.05% concentration. At 0.1% concentration, the subculture results were consistent with the above drug exposure results. We did not find any regrowth in samples of three top hits, oregano, cinnamon bark and clove bud (Figure 3Ab-d). However, citronella, geranium bourbon and wintergreen could not completely kill the stationary phase *B. burgdorferi* with many spirochetes being visible after 21-day subculture (Figure 3Ae-g). Subculture also confirmed the activity of carvacrol by showing no spirochete regrowth in the 0.1% carvacrol treated samples. In p-cymene and α-terpinene subculture samples, we observed growth even in 0.1% concentration samples. At 0.05% concentration, we observed no spirochetal regrowth after 21-day subculture in the oregano and cinnamon bark treated samples (Figure 3Bb, c), despite some very tiny aggregated microcolonies were found after treatment (Figure 2Bb, c). Although the clove bud showed better activity than the cinnamon bark at 0.05% concentration (Table 2), interestingly, clove bud could not sterilize the *B. burgdorferi* stationary phase culture, as they all had visible spirochetes growing after 21-day subculture (Figure 3Bc, d). Additionally, 0.05% citronella, geranium bourbon and wintergreen could not kill all *B. burgdorferi* since many viable spirochetes were observed in the 21-day subculture (Figure 3Be-g). Remarkably, 0.05% carvacrol sterilized the *B. burgdorferi* stationary phase culture as shown by no regrowth after 21-day subculture (Figure 3Bh).

**FIG 3.**
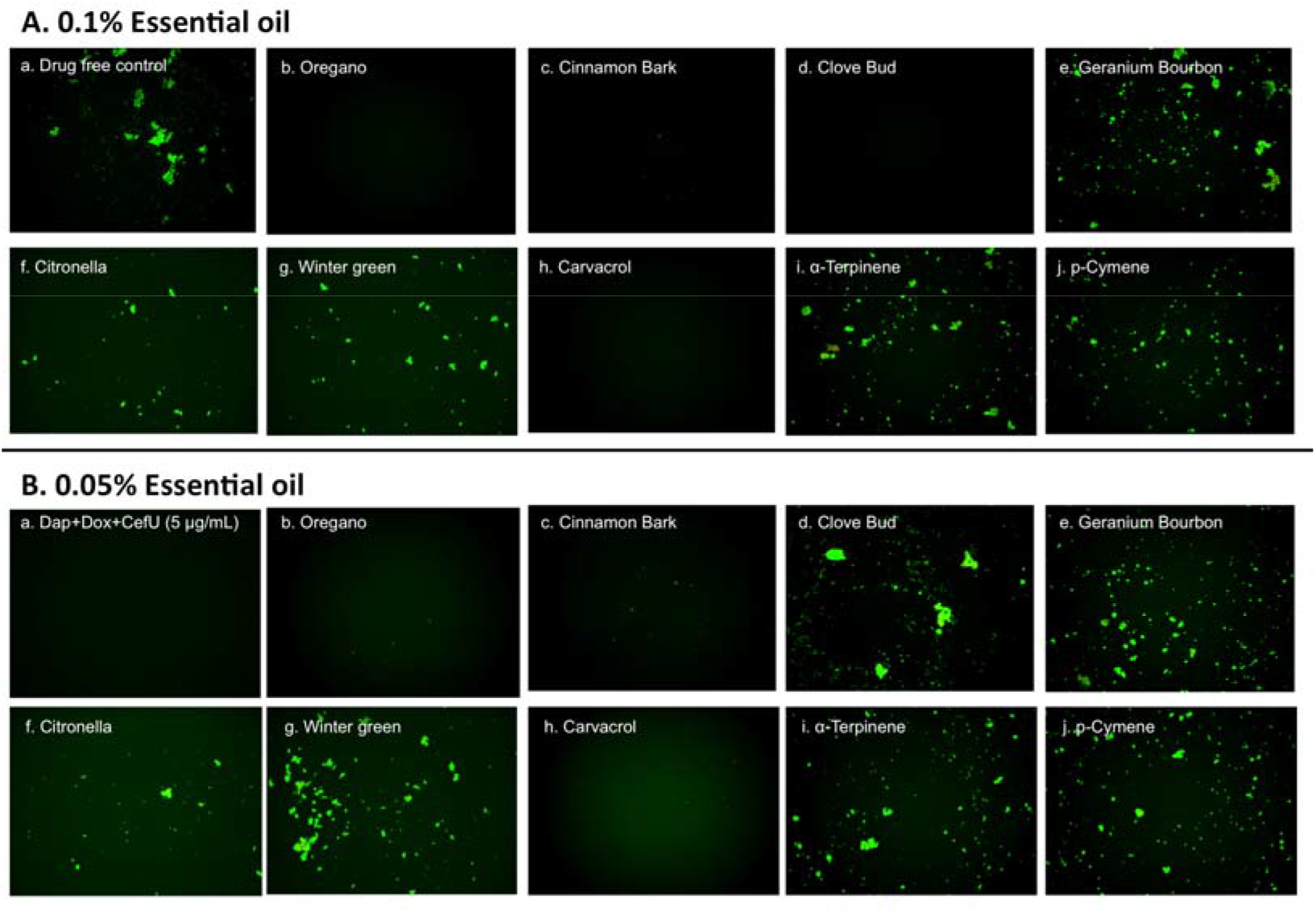
Subculture of *B. burgdorferi* after treatment with essential oils. A *B. burgdorferi* stationary phase culture (7-day old) was treated with the indicated essential oils at 0.1% (A) or 0.05% (B) for 7 days followed by washing and resuspension in fresh BSK-H medium and subculture for 21 days. The viability of the subculture was examined by SYBR Green I/PI stain and fluorescence microscopy.

## DISCUSSION

Previous *in vitro* studies showed that certain essential oils have bacteriostatic and/or bactericidal activity against on multidrug resistant Gram-negative clinical isolates (23). In this study, we tested 34 essential oils from different plants on non-growing stationary phase *B. burgdorferi* as a model of persister drug screens. We were able to identify 23 essential oils that are more active than 40 μM daptomycin at 1% concentration, 3 of which, i.e. oregano, clove bud and cinnamon bark, highlighted themselves as having a remarkable activity even at a very low concentration of 0.125% (Table 1). Among them oregano and cinnamon bark essential oil had the best activity as shown by completely eradicating *B. burgdorferi* even at 0.05% concentration. In a previous study, oregano essential oil was found to have antibacterial activity against Gram-positive and Gram-negative bacteria (22). Here, for the first time, we identified oregano essential oil as having a highly potent activity against stationary phase *B. burgdorferi*. We tested three major ingredients of oregano essential oil (carvacrol, p-cymene and α-terpinene) on *B. burgdorferi*, and found carvacrol is the major active component, which showed similar activity as the complete oregano essential oil (Figures 2 and 3). In addition, we noted that oregano essential oil can dramatically reduce the size of aggregated biofilm-like microcolonies compared to the antibiotic controls (Figure 1). After treatment with 0.25% oregano essential oil, only some dispersed tiny red aggregated cells were left in the culture (Figure 1C). Interestingly, we observed that amount and size of aggregated biofilm-like microcolonies of *B. burgdorferi* dramatically reduced with increasing concentrations of oregano oil, as aggregated biofilm-like structures vanished after treatment with 0.5% or 1% oregano essential oil. When we reduced the concentration of oregano essential oil to 0.05%, it could not eradicate stationary phase *B. burgdorferi* (residual viability 56%, Figure 2Bb) but the size of aggregated microcolonies decreased significantly. By contrast, daptomycin could kill the aggregated biofilm-like microcolonies of *B. burgdorferi* as shown by red aggregated microcolonies but could not break up the aggregated microcolonies even at the highest concentration of 40 μM (Figure 1A). It has been shownthat carvacrol and other active compositions of oregano essential oil could disrupt microbial cell membrane (19). Future studies are needed to determine whether oregano essential oil and other active essential oils have similar membrane disruption activity and could destroy the aggregated biofilm structures of *B. burgdorferi*.

We also noted that some essential oils such as oregano and cinnamon bark had relatively high residual viability percentage (Table 2) at low concentration of 0.05% but their treated *B. burgdorferi* cells did not grow in the subculture study (Table 2; Figure 3Bb, c). We speculate that these essential oils could dissolve the dead *B. burgdorferi* cells presumably due to their high lipophilicity. The reduction of number of dead red cells by the essential oil made the residual viability percentage increase, although the amount of live cells obviously decreased as well (Figure 2Ab-d, Bb-c). In addition, these essential oils may also permanently damage or inhibit the growth of *B. burgdorferi* during the treatment, such that even in the fresh medium, the residual *B. burgdorferi* cells still could not regrow.

Meanwhile, we found that at a high concentration (above 1%) lemongrass or oregano essential oil showed apparent high residual viability percentage by the SYBR Green I/PI plate assay, compared with the microscopy counting data (Table 1, Figure 1A). This may be caused by strong autofluorescence of these essential oils that severely interfere with the SYBR Green I/PI assay. We studied the emission spectral of lemongrass essential oil using Synergy H1 multi-mode reader and found lemongrass essential oil emits the strongest autofluorescence. The peak fluorescence of lemongrass essential oil is at 520 nm that overlaps with the green fluorescence of SYBR Green I dye (peak is at 535 nm). The strong autofluorescence caused the abnormal residual viability percentage (above 100% in Table 1) using SYBR Green I/PI plate assay. We also found oregano essential oil emits autofluorescence at 535 nm, which pushed the green/red fluorescence ratio higher than their true values (Table 1). However, we were able to solve this problem by using fluorescence microscopy as a more reliable measure to confirm the results of SYBR Green I/PI plate reader assay (13, 21).

Additionally, we found cinnamon bark and clove bud essential oils showed excellent activity against *B. burgdorferi*. Cinnamon bark essential oil eradicated the stationary phase *B. burgdorferi* even at 0.05% concentration (Table 2) while clove bud essential oil showed sterilization at 0.1% or above concentration. Extractions of cinnamon bark and clove bud have been used as flavors for food processing. Based on this discovery, effective oral regimens with low side effect may be developed to fight against Lyme disease in future studies.

In a previous study, it has been found that volatile oil from *Cistus creticus* showed growth inhibiting activity against *B. burgdorferi in vitro* (24) but its activity against stationary phase bacteria enriched in persisters was not evaluated. In this study, we tested six *Citrus* plants *(Citrus bergamia, Citrus sinensis, Citrus limonum, Citrus aurantifolia, Citrus racemosa, Citrus reticulata)* on the stationary phase *B. burgdorferi* culture and found bergamot *(Citrus bergamia)* had high activity (residual viability 12%) at 1% concentration but the other *Citrus* essential oils did not show good activity against *B. burgdorferi* compared with clinically used doxycycline, cefuroxime or ciprofloxacin (Table 1).

Although we found several essential oils (oregano, cinnamon bark, clove bud) that have excellent sterilizing activity against *B. burgdorferi* stationary phase cells in vitro (Table 1), the effective dose that will show equivalent activity in vivo is unknown at this time largely because the active ingredients in the active essential oils and the pharmacokinetic profile of the active ingredients are not all known. Future studies are needed to identify the active ingredients of the active essential oils and determine their effective dosage in vivo. Identification of active components or active component combinations from essential oils may help to eliminate the quality difference of natural products. However, we were able to identify carvacrol as the most active ingredient in oregano essential oil, and its pharmacokinetics has been studied as a feed addition in pigs (25) and topical oil in cattle (26). In the rat model, the calculated LD50 of carvacrol is 471.2 mg/kg (27). We noticed that the 0.05% of carvacrol used here, which is equivalent to 0.48 μg/mL or 3.2 μM and completely eradicated *B. burgdorferi* stationary phase cells in subculture (Figure 3), is lower than the peak plasma concentration (3.65 μg/mL) in the swine study (25). These findings favor the application of carvacrol in future treatment studies. Importantly, carvacrol seems to be more active than daptomycin, the most active persister drugs against *B. burgdorferi* (13, 14). In this study, 0.1% carvacrol (6.4 μM) showed much higher activity (2% residual viability) than 5 μM daptomycin (45% residual viability) (Table 1 and 2). In addition, 0.05% carvacrol (3.2 μM) could eradicate *B. burgdorferi* stationary phase cells with no regrowth in subculture, but 10 μg/mL daptomycin (6.2 μM), by contrast, could not completely kill *B. burgdorferi* stationary phase cells as shown by regrowth in subculture (14). Furthermore, carvacrol showed remarkable activity against not only stationary phase *B. burgdorferi* but also log phase replicating cells with very low MIC (0.16-0.31 μg/mL). However, there is limited safety information on carvacrol in humans. In mice, carvacrol has been given at 40 mg/kg daily for 20 days with no apparent toxicity (28). However, carvacrol and other active components of essential oil showed certain cytotoxicity (IC50 of carvacrol was 200-425 μM) (29, 30) on mammalian cells and genotoxic activity *in vivo* (even the lowest dose of 10 mg/kg) (31). In addition, it is well known that some effective drugs identified in vitro may fail when tested *in vivo*. Thus, adequate animal studies are needed to confirm the safety and efficacy of the active essential oils in *in vivo* setting before human studies.

In summary, we found that many essential oils had varying degrees of activity against stationary phase *B. burgdorferi*. The most active essential oils are oregano, cinnamon bark, and clove bud, which seem to have even higher activity than the persister drug daptomycin. A particularly interesting observation is that these highly active essential oils had remarkable biofilm-dissolving capability and completely eradicated all stationary phase cells with no regrowth. In addition, carvacrol was found to be the most active ingredient of oregano with high activity against *B. burgdorferi* stationary phase cells. Future studies are needed to test whether carvacrol could replace the persister drug daptomycin in drug combinations against more resistant biofilm-like structures and for treating persistent borrelia infections in animal models and in patients.

## MATERIALS AND METHODS

### Strain, media and culture techniques

Low passaged (less than 8 passages) *B. burgdorferi* strain B31 5A19 was kindly provided by Dr. Monica Embers (15). The *B. burgdorferi* B31 strain was grown in BSK-H medium (HiMedia Laboratories Pvt. Ltd.) and supplemented with 6% rabbit serum (Sigma-Aldrich, St. Louis, MO, USA). All culture medium was filter-sterilized by 0.2 μm filter. Cultures were incubated in sterile 50 ml conical tubes (BD Biosciences, California, USA) in microaerophilic incubator (33°C, 5% CO_2_) without antibiotics. After incubation for 7 days, 1 ml stationary-phase *B. burgdorferi* culture (~10^7^ spirochetes/mL) was transferred into a 96-well plate for evaluation of potential anti-persister activity of essential oils (see below).

### Essential oils and drugs

A panel of essential oils was purchased from Plant Therapy (ID, USA), Natural Acres (MO, USA), or Plant Guru (NJ, USA). Carvacrol, p-cymene, and α-terpinene were purchased from Sigma-Aldrich (USA). Essential oils were added to *B. burgdorferi* cultures to form aqueous suspension by vortex. Immediately the essential oil aqueous suspension was serially diluted to desired concentrations followed by addition to *B. burgdorferi* cultures. Doxycycline (Dox), cefuroxime (CefU), (Sigma-Aldrich, USA) and daptomycin (Dap) (AK Scientific, Inc, USA) were dissolved in suitable solvents (32, 33) to form 5 mg/ml stock solutions. The antibiotic stocks were filter-sterilized by 0.2 μm filter and stored at −20°C.

### Microscopy

The *B. burgdorferi* cultures were examined using BZ-X710 All-in-One fluorescence microscope (KEYENCE, Inc.). The SYBR Green I/PI viability assay was performed to assess the bacterial viability using the ratio of green/red fluorescence to determine the live:dead cell ratio, respectively, as described previously (13, 34). This residual cell viability reading was confirmed by analyzing three representative images of the bacterial culture using epifluorescence microscopy. BZ-X Analyzer and Image Pro-Plus software were used to quantitatively determine the fluorescence intensity.

### Evaluation of essential oils for their activities against *B. burgdorferi* stationary phase cultures

To evaluate the essential oils for possible activity against stationary phase *B. burgdorferi*, aliquots of the essential oils or drugs were added to 96-well plate containing 100 μL of the 7-day old stationary phase *B. burgdorferi* culture to obtain the desired concentrations. In the primary essential oil screen, each essential oil was assayed in four concentrations, 1%, 0.5%, 0.25% and 0.125% (v/v) in 96-well plate. The active hits were further confirmed with lower 0.1% and 0.05% concentration; all tests were run in triplicate. All the plates were incubated at 33°C and 5% CO2 without shaking for 7 days when the residual viable cells remaining were measured using the SYBR Green I/PI viability assay and epifluorescence microscopy as described (13, 34).

### Antibiotic susceptibility testing

To qualitatively determine the effect of essential oils in a high-throughput manner, 10 μl of each essential oil from the pre-diluted stock was added to 7-day old stationary phase *B. burgdorferi* culture in the 96-well plate. Plates were sealed and placed in 33°C incubator for 7 days when the SYBR Green I/ PI viability assay was used to assess the live and dead cells as described (13). Briefly, 10 μl of SYBR Green I (10,000 × stock, Invitrogen) was mixed with 30 μl propidium iodide (PI, 20 mM, Sigma) into 1.0 ml of sterile dH2O. Then 10 μl staining mixture was added to each well and mixed thoroughly. The plates were incubated at room temperature in the dark for 15 minutes followed by plate reading at excitation wavelength at 485 nm and the fluorescence intensity at 535 nm (green emission) and 635 nm (red emission) in microplate reader (HTS 7000 plus Bio Assay Reader, PerkinElmer Inc., USA). With least-square fitting analysis, the regression equation and regression curve of the relationship between percentage of live and dead bacteria as shown in green/red fluorescence ratios was obtained. The regression equation was used to calculate the percentage of live cells in each well of the 96-well plate.

The standard microdilution method was used to determine the MIC of carvacrol, based on inhibition of visible growth of *B. burgdorferi* by microscopy. Carvacrol was added to *B. burgdorferi* cultures (1 × 10^4^ spirochetes/mL) to form aqueous suspension by vortex. The carvacrol suspension was two-fold diluted from 0.5% (4.88 μg/mL) to 0.008% (0.08 μg/mL). All experiments were run in triplicate. *B. burgdorferi* culture was incubated in 96-well microplate at 33 °C for 7 days. Cell proliferation was assessed using the SYBR Green I/PI assay and BZ-X710 All-in-One fluorescence microscope (KEYENCE, Inc.).

### Subculture studies to assess viability of the of essential oil-treated *B. burgdorferi* organisms

A 7-day old *B. burgdorferi* stationary phase culture (500 μl) was treated with essential oils or control drugs for 7 days in 1.5 ml Eppendorf tubes as described previously (14). After incubation at 33 °C for 7 days without shaking, the cells were collected by centrifugation and rinsed with 1 ml fresh BSK-H medium followed by resuspension in 500 μl fresh BSK-H medium without antibiotics. Then 50 μl of cell suspension was transferred to 1 ml fresh BSK-H medium for subculture at 33 °C for 20 days. Cell proliferation was assessed using SYBR Green I/PI assay and epifluorescence microscopy as described above.

## ACKNOWLEDGMENTS

We acknowledge the support of this work by Steven & Alexandra Cohen Foundation, Global Lyme Alliance, Lyme Disease Association, NatCapLyme, and Steve Sim Fund. YZ was supported in part by NIH grants AI099512 and AI108535.

